## supplemental figures for "The non-mitotic role of HMMR in regulating the localization of TPX2 and the dynamics of microtubules in neurons"

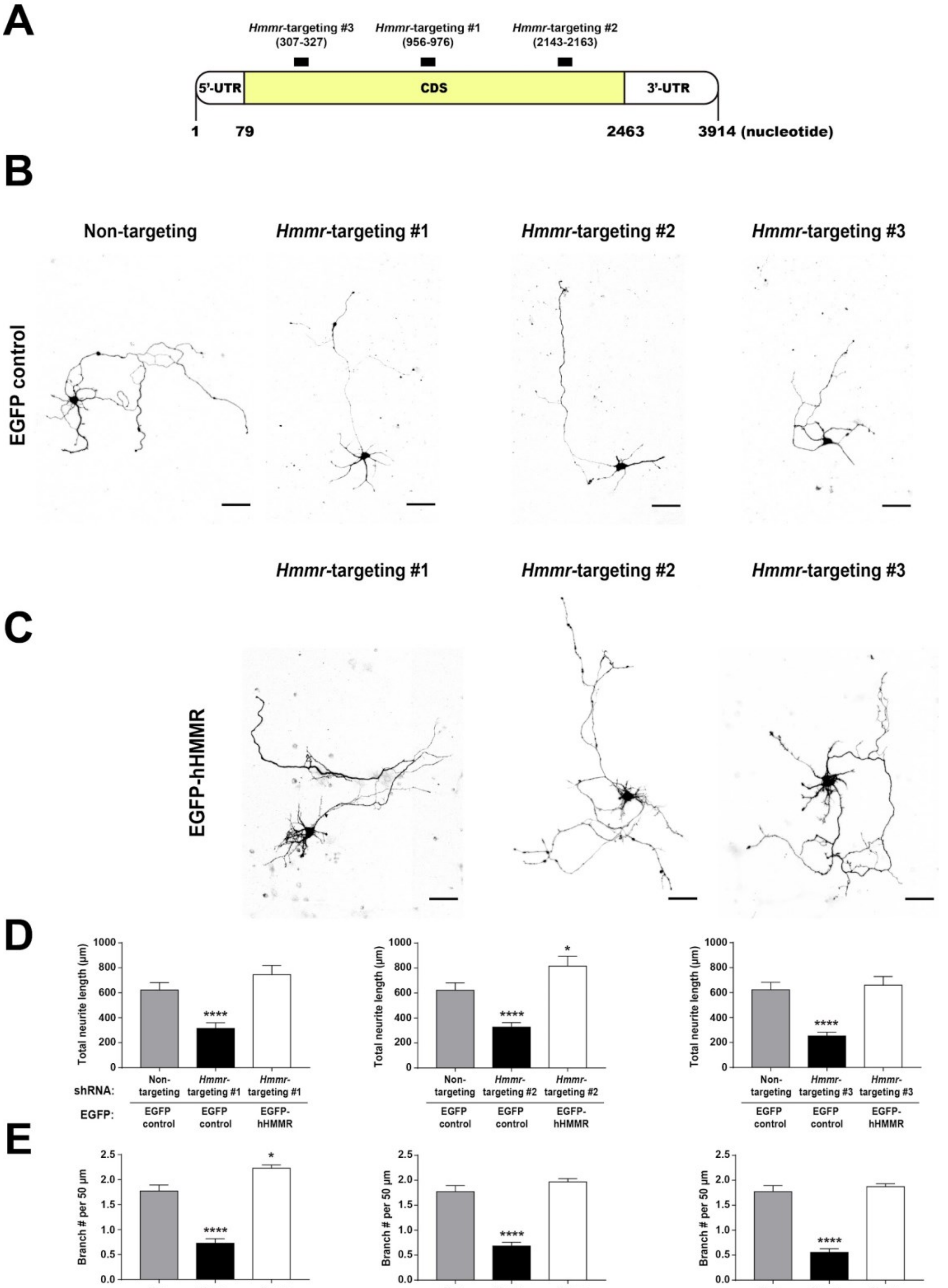


**Figure S1. Overexpressing human HMMR rescues the effect of HMMR depletion in mouse neurons.**

(A) Schematic representation of shRNA targeting regions of the mouse *Hmmr* mRNA. The untranslated regions (UTR) are shown in white and the coding sequence (CDS) in yellow. Black bars indicate shRNA targeting regions and the mRNA is numbered by the nucleotide sequence. Representative EGFP images of mouse hippocampal neurons co-transfected with the indicated shRNA- and the control cytosolic EGFP- (B) or EGFP-hHMMR (C) expressing plasmids on 2 DIV and fixed on 5 DIV. Images are inverted to improve visualization. The scale bars present 50 µm. Quantification of (D) total neurite length per neuron and (E) normalized branch density (branch number in 50 µm of neurite) for neurons shown in panel B-C. * $P<0.05$; **** $P<0.0001$, one-way ANOVA followed by Dunnett's post-hoc test. All bar graphs are expressed as mean ± s.e.m. from three independent repetitions. More than 50 neurons were analyzed per condition per repeat.

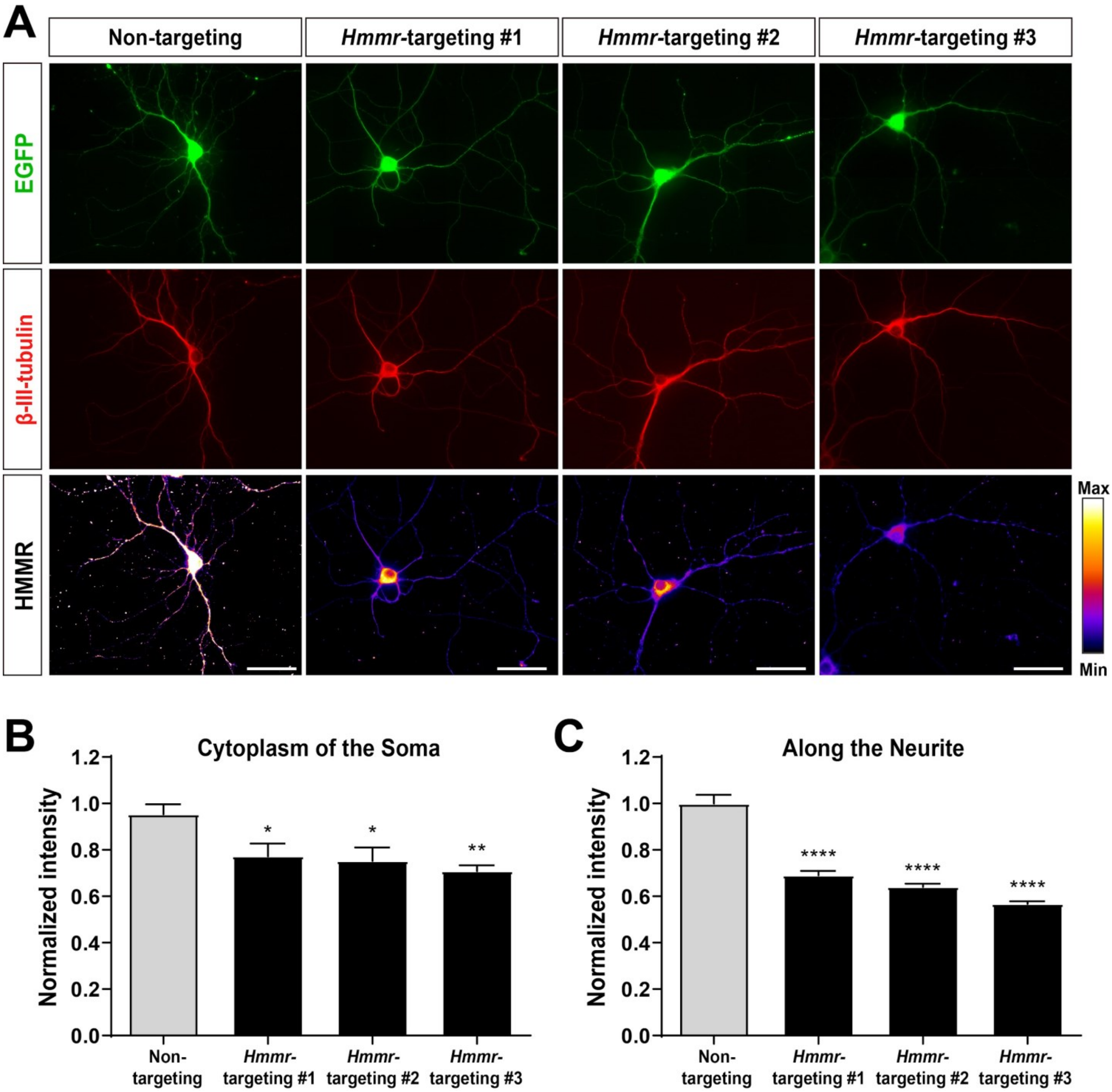


**Figure S2. Validation of the HMMR antibody.**

(A) Representative images of 10 DIV mouse hippocampal neurons co-expressing the indicated *Hmmr*-targeting shRNA and EGFP. Neurons were immunofluorescence stained with antibodies against β-III-tubulin (red) and HMMR (pseudocolor). All scale bars present 50 μm. Only neurons possessing both β-III-tubulin and EGFP signals were quantified. Quantification of HMMR intensity in the soma (B) and along the neurite (C). * $P<0.05$; ** $P<0.01$; **** $P<0.0001$, one-way ANOVA followed by Dunnett's post-hoc tests. Both bar graphs are expressed as mean ± s.e.m. from three independent repetitions. More than 30 neurons were analyzed per condition per repeat.

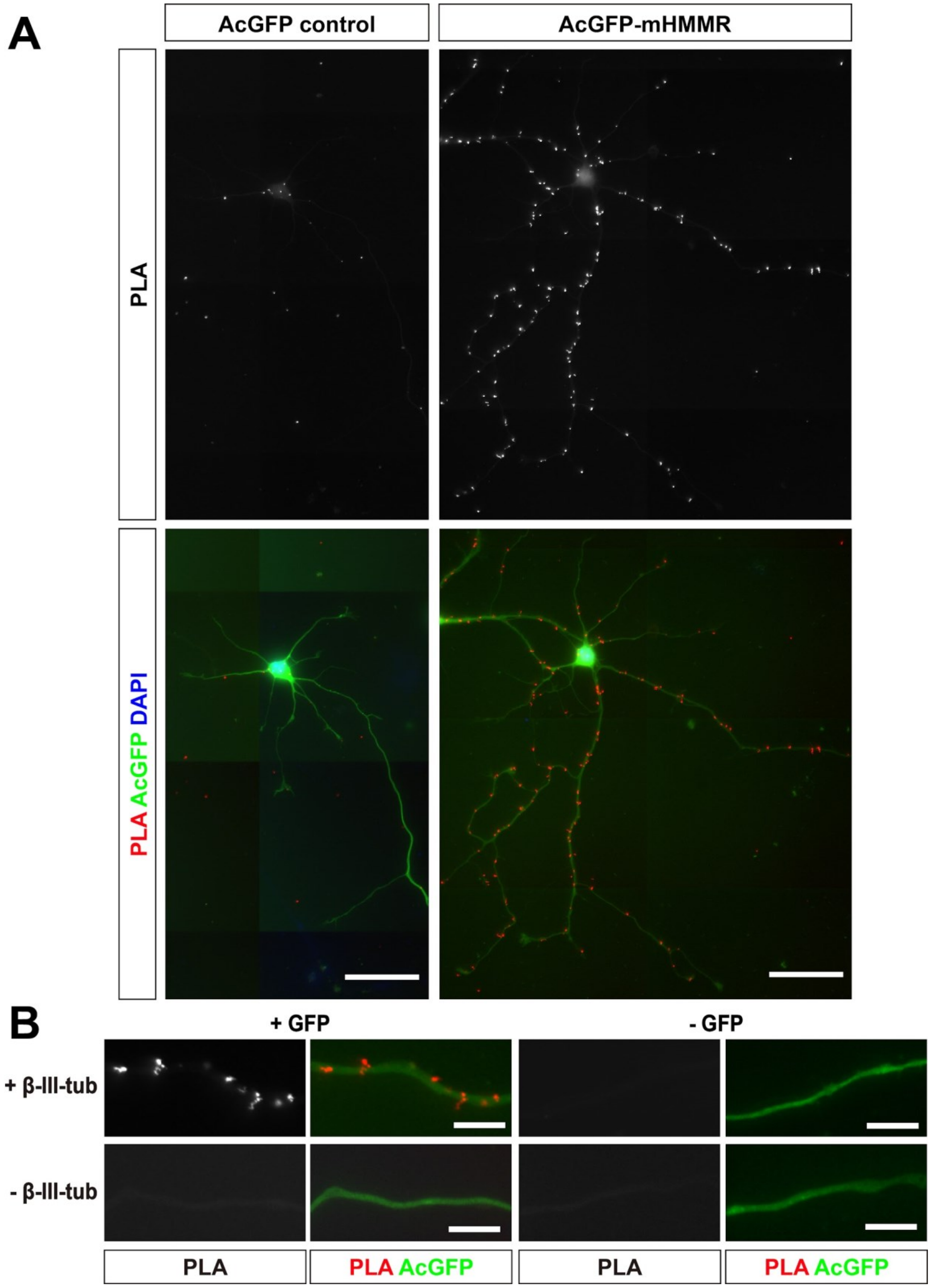


**Figure S3. Transiently expressed AcGFP-mHMMR associates with microtubules in neurons.**

Dissociated hippocampal neurons were transfected with AcGFP control (left) or AcGFP-mHMMR (right) on 0 DIV and fixed on 7 DIV. (A) PLA images of AcGFP control (upper left) or AcGFP-mHMMR (upper right) and β-III-tubulin in 7DIV dissociated hippocampal neurons. Nuclei were visualized using DAPI (blue) and the general appearance of neurons was visualized using the AcGFP signal (green). All images have the same scale and the scale bars present 50 µm. (B) PLA puncta were present along the neurite shaft only when antibodies against AcGFP and β-III-tubulin were both present. All images have the same scale and the scale bars represent 10 µm.

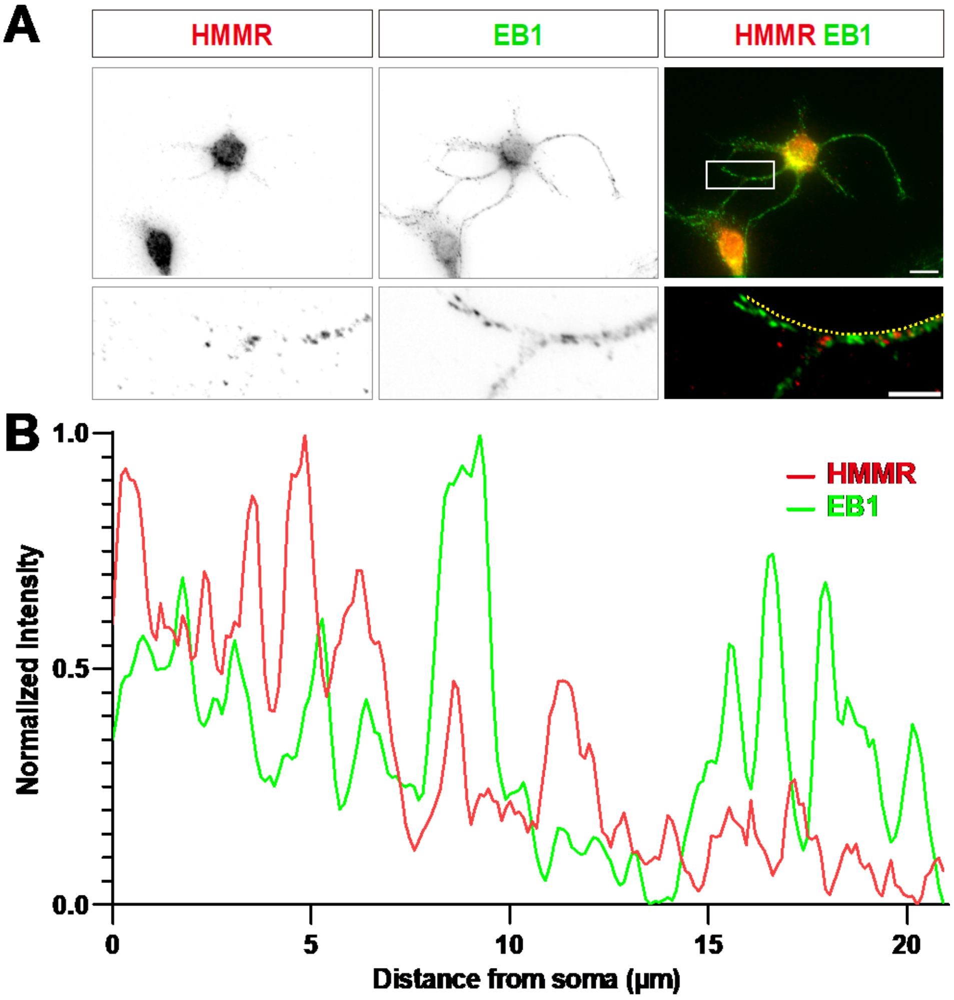


**Figure S4. HMMR does not colocalize with microtubule plus-ends in neurons.**

(A) Representative images of a 3 DIV hippocampal neuron immunofluorescence stained with antibodies against HMMR and EB1. The white box in the upper right panel indicates the magnified region shown in lower panels. The scale bars in the upper and lower panels represent 10 μm and 5 μm, respectively. (B) The linescan along the yellow dotted line in the lower right of panel A. The signal intensity of HMMR and EB1 is shown in red and green, respectively. The average Pearson correlation coefficient is 0.28±0.25. More than 300 neurites from three independent repeats were quantified.
